# Using Tools as Cues for Motor Adaptation in Virtual Reality

**DOI:** 10.1101/2025.09.02.673800

**Authors:** Andrew Michael King, Jacob J. Boulrice, Shanaathanan Modchalingam, Laura Mikula, Bernard Marius ‘t Hart, Denise Y P Henriques

## Abstract

Humans are adept at using multiple tools, often switching between them even when each involves distinct and potentially conflicting motor demands. This study investigated how different features of a tool influence the formation of distinct motor memories during dual adaptation to opposing visuomotor perturbations. Using an immersive virtual reality (VR) setup, participants performed an aiming task in which they launched a ball toward a target using virtual tools, each consistently associated with a different visuomotor rotation. We manipulated tool features across three groups: one in which tools differed only in colour (Colour Control), one in which they differed in shape but involved similar movement patterns (Motor Congruent), and one in which they differed both in shape and in how they were operated—for example, a paddle-like forward swing to propel the ball versus a draw-and-release motion similar to a slingshot (Motor Incongruent). A fourth control group that adapted to a single perturbation with a single tool at a time was also included. Only the Motor Incongruent group demonstrated robust dual adaptation and clear aftereffects, comparable to those observed during single-tool learning. These results suggest that distinct modes of tool operation play a critical role in supporting the formation and retention of separate internal models during sensorimotor adaptation.

**New & Noteworthy:** Humans often switch between tools with conflicting motor demands. Using immersive virtual reality, we tested whether visual features or operational differences support dual adaptation to opposing visuomotor perturbations. Only when tools differed in both shape and mode of operation did participants show robust adaptation and aftereffects, comparable to single-tool learning. These findings highlight the critical role of movement-relevant cues in forming distinct motor memories for flexible tool use.

## Introduction

Humans excel at using a variety of tools, often switching fluidly between them—even when they require opposing motor actions. This flexibility suggests that the motor system can form and retrieve distinct internal models to govern tool-specific actions without interference. However, the conditions under which such separate motor memories can be acquired and maintained remain unclear.

Sensorimotor adaptation studies have shown that learning a novel mapping between motor commands and their expected sensory outcomes—such as adapting to a rotated visual cursor—relies on sensory prediction errors that update internal models of movement ^1–4^. However, when individuals are required to adapt to two opposing perturbations within the same session, performance often suffers due to interference. In this context, interference refers to the tendency for learning one perturbation (e.g., a clockwise rotation) to partially overwrite, disrupt, or destabilize the motor memory formed for the opposing perturbation (e.g., a counterclockwise rotation). As a result, the motor system fails to maintain two distinct internal models, leading to reduced retention, slower learning, or a blending of the two learned states into a compromised intermediate solution. This limitation suggests that, without additional contextual cues to signal when each perturbation should be applied, the nervous system treats the conflicting experiences as arising from a single task environment and updates a single control policy. By contrast, when clear contextual cues are provided—such as distinct visual tools, postures, or task rules—the motor system can assign each perturbation to a separate internal model, thereby reducing interference and enabling robust dual adaptation ^5–14^.

Prior work has demonstrated that intrinsic contextual cues—those tied to the motor execution itself, such as movement direction, posture, or lead-in movements— support dual adaptation by recruiting distinct control policies ^5,6,11–21^In contrast, extrinsic cues—such as color or tool shape—often fail to enable interference-free learning unless made task-relevant or tightly associated with specific sensorimotor demands ^15,17,22,23^.

Tools present a particularly interesting case, as they combine both extrinsic visual features and intrinsic motor properties. Some studies have shown that tools can support dual adaptation when they differ in their control dynamics or when paired with distinct contact goals ^9,10,12,24^. However, the relative importance of visual versus motoric features in supporting the formation of separate internal models remains unresolved.

In this study, we investigated whether visual-identity and operational features of tools can serve as effective contextual cues for dual adaptation in a visuomotor rotation task. Using immersive virtual reality (VR), participants adapted to two opposing rotations that altered the direction of ball motion, with each rotation consistently paired with a distinct tool. Importantly, each tool launched the ball in a unique manner, requiring participants to engage with it using different movement dynamics—for example, one tool involved a smooth paddle-like swing to propel the ball, whereas another required a draw-and-release action similar to a slingshot. Across experimental groups, we systematically manipulated whether these tool pairs differed only in their visual shape (providing purely extrinsic cues), in both shape and the specific movement required to operate them (combining extrinsic and intrinsic cues), or solely in color (a minimal extrinsic cue without functional differences). By doing so, our goal was to determine whether movement-specific operational properties are necessary to support robust dual adaptation, or whether visual features alone can serve as sufficient contextual signals in the absence of distinct motor requirements.

## Methods

### Participants

A total of 144 right-handed undergraduate students (98 female, mean age = 21.98 ± 4.62 years) participated in the study. Participants belonged to one of four groups: Motor Incongruent (n = 40), Motor Congruent (n = 40), Colour Control (n = 44), and Single Adaptation (n = 20). All had normal or corrected-to-normal vision and provided informed consent. Participants received course credit for participation. The study protocol was approved by York University’s Human Participants Review Sub-committee.

### Apparatus and Virtual Environment

Participants were seated at a table and wore an Oculus Rift CV1 virtual reality headset (resolution: 1080 × 1200 pixels per eye; refresh rate: 90 Hz). Head and hand movements were tracked using three external Oculus sensors, which sampled at 60 Hz with sub-millimetre precision (<1 mm). Participants used a hand-held Oculus Touch controller to interact with objects in a custom-built virtual environment developed in Unity. The virtual scene was designed to match the physical dimensions of the table (3.5 m wide × 2 m deep) to provide consistent spatial alignment between real and virtual space (as with Modchalingam et al., 2024, 2025). Trial presentation and data collection were managed using the Unity Experimenter Framework^26^, which supported synchronized stimulus control and response logging throughout the experiment.

### Task and Stimuli

Participants performed a tool-mediated ball-launching task. On each trial, they used one of three virtual tools—a paddle, slingshot, or curling rod (Fig 1C)—to launch a virtual ball (5 cm diameter) toward a target (5 cm) positioned 85 cm away in one of four directions (75°, 85°, 95°, 105° polar angle, Fig 1A). Tools differed in shape (Fig 1C-D) and pre-launch movement (Fig 1D, 4B): the paddle and curling rod involved a forward strike, while the slingshot required a backward pull and release. Each Group only trained with two of the three tools, were used to manipulate motor congruency between tools (as per Fig 1D).

**Figure 1.**
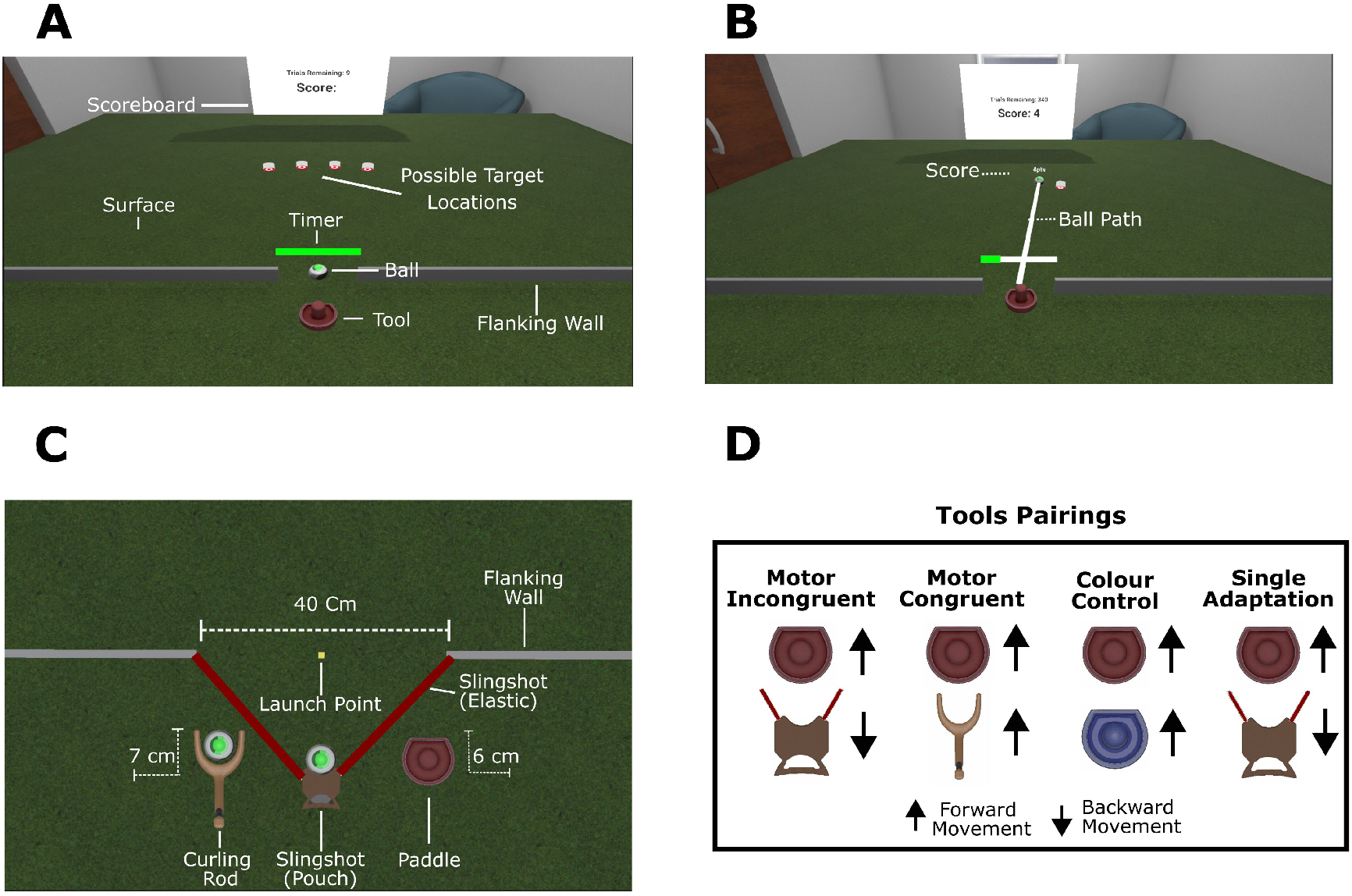
Experimental apparatus. A) Virtual environment in the VR experiment at the beginning of each trial. Once participants grabbed the tool, the timer would begin to deplete, prompting them to complete their shot toward the target. All four potential target locations — 75°, 85°, 95°, and 105° — are shown. B) The virtual environment in the VR experiment at the end of each trial. Participants score and trials remaining would be displayed on the scoreboard. In this example, the 75° target is shown. C) Rendering of the tools in the virtual environment. D) Illustration of the tool pairs used in each group. Arrows indicate whether a forward or backward movement was required to shoot the ball. Each Dual Adaptation group (Motor Incongruent, Motor Congruent, Colour Control) trained with two opposing rotations, each paired with a distinct tool. For the Single Adaptation groups (shown on the right), only one tool and one rotation direction were used/presented for a long training block.

Ball trajectories after launch was governed by Unity’s physics engine, PhysX, with a maximum launch-velocity cap of 22 cm/s. Flanking walls on the virtual table ensured that the ball was released from the same restricted area for all tools, and the tool positions were constrained so they could not extend beyond this launch zone At the end of each trial, participants received visual feedback showing the ball’s trajectory and a performance score (0-10 points), based on the minimum distance between the ball and the target (e.g., Fig 1B). A green timer bar (Fig 1A-B) prompted participants to launch the ball within 1 s (3 s during practice). Late responses or shots deviating more than 20 cm from the target received a score of 0.

### Experimental Design

The experiment employed a between-subjects design with four participant groups, see Fig 1D. Three groups completed a dual adaptation (DA) condition, in which each participant used two distinct tools consistently associated with opposing visuomotor perturbations (clockwise [CW] or counterclockwise [CCW]). The Motor Congruent group trained with the paddle and curling rod, which required similar forward launching movements. The Motor Incongruent group trained with the paddle and slingshot, which required contrasting movement patterns (forward vs. backward), introducing greater sensorimotor dissimilarity between tools. The Colour Control group used two visually distinct paddles (red and blue) that differed only in colour and not in movement dynamics, allowing us to isolate the role of visual cues alone. A fourth group, the Single Adaptation (SA) group, served as a control group and completed two separate adaptation blocks with a single tool and perturbation pairing per block, with no interleaving of perturbation.

All participants completed four experimental phases: a Practice phase, a Baseline phase, a Training phase, and a Washout phase, as illustrated in Fig 2A. The Practice phase familiarized participants with the mechanics of the ball-launching task using unperturbed feedback (24 trials per tool). The Baseline phase consisted of 80 trials with veridical feedback (40 per tool) to assess initial performance (illustrated in Fig 2B, top row and Fig 2C, left panel). In the Training phase, a 30° rotation (CW or CCW) was imposed on the ball trajectory (illustrated in Fig 2B, bottom row and Fig 2C, right panel). In the dual adaptation groups (Fig 2A, top row), each rotation was consistently paired with one of the two tools, and tools alternated every 8 trials in mini-blocks throughout the phase (160 trials total; 80 per tool). Each block of these 8 trials is represented by a single rectangle in Fig 2A, and with different shades of same colour associated with different tools. In contrast, participants in the Single Adaptation group (Fig 2A, bottom row) completed 80 consecutive trials with one tool and rotation before switching to the other tool and opposing perturbation for an additional 80 trials. To compensate for the visuomotor rotation introduced during the Training phase, participants had to adjust their hand movements in the direction opposite to the imposed rotation in order to accurately aim the ball at the target. The order of tool and perturbation was counterbalanced across participants. During the final Washout phase, participants completed 48 aligned trials (24 per tool) —with veridical feedback identical to Baseline— to assess aftereffects and de-adaptation. Background textures differed between aligned and rotated trials (Fig 2B), but participants were not informed of these changes, nor were they explicitly told about the visuomotor rotations. Short rest breaks were given between phases, and participants kept their headsets on throughout the session.

**Figure 2.**
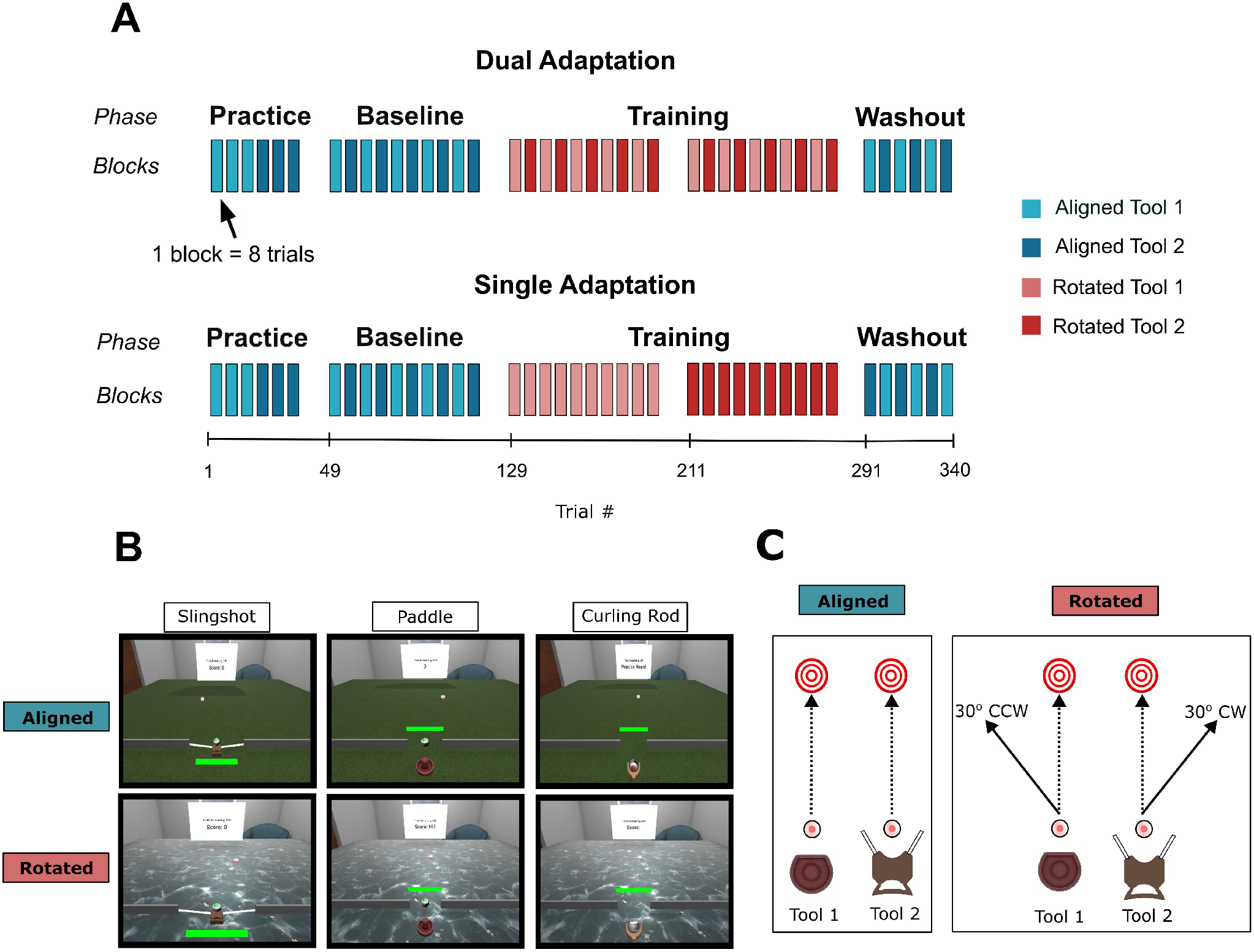
Layout of the experimental design. A) Participants in all groups completed 4 different experimental phases. Every vertical bar represents one block of 8 tool-specific trials. In Baseline and Washout phases, no perturbation was imposed on the ball (aligned trials). The Training phase involved a 30° angular perturbation on the ball’s movement direction (rotated trials). In the Dual Adaptation conditions, participants alternated between tools every block. In the Single Adaptation condition, participants only alternated tools in the Baseline and Washout phases. During Training, these participants used one tool for 80 consecutive trials at a time. Tool order and rotation direction were counterbalanced across all participants. B) Visualization of aligned and rotated trials for the slingshot, paddle, and curling rod trials. C) Illustration of different trial types. In aligned trials, the ball rolls forward in the intended aiming direction, while in rotated trials the ball travels either 30° CW or CCW depending on the tool used.

### Procedure

The participant’s task was to use the provided tool to shoot a virtual ball toward a visible target. To initiate each trial, participants moved the VR hand-held controller to the tool’s location. As the controller and hand had no visible representation aside from the tool itself, the controller vibrated when it came within 1 cm of the tool to provide haptic confirmation of proximity. Upon reaching the tool, participants pressed and held the trigger button to “grab” it, which initiated a brief countdown signaled by a green horizontal bar displayed on the screen (Fig 1A–B). This countdown lasted one second during experimental trials and three seconds during practice trials, after which participants were prompted to quickly and accurately shoot the ball toward the target. Shooting the ball involved intuitive, tool-specific movements; for example, the paddle and rod required a forward push, while the slingshot required a backward pulling motion.

Each trial ended when the ball either hit the target, slowed to a speed below 1 cm/s, or reached a radial distance of 120 cm from the launch point. At that point, a white line appeared to indicate the ball’s path, providing participants with visual feedback on their accuracy (Fig 1B). To motivate participants, a score between 0 and 10 was awarded on each trial based on their minimum error—that is, the smallest distance between the ball and the target during the trajectory. An additional 5 bonus points were awarded if the ball hit the center of the target. The cumulative score and number of remaining trials were displayed throughout the session on an in-VR scoreboard (Fig 1A–B). Trials in which the ball missed the target by more than 20 cm or was launched after the countdown expired received a score of 0.

### Data Analysis

This study investigated how tool identity (extrinsic cues) and tool operation (intrinsic cues) contribute to the concurrent learning of tool-pair associations, each linked to different visuomotor perturbations. By manipulating the congruency between the movements required to operate each tool and its visual identity, we evaluated whether these characteristics support the formation of separate motor memories under conditions of dual adaptation.

The primary dependent variable was angular error, defined as the difference between the direction of the ball’s trajectory and the target direction. To allow comparisons across the two opposing perturbation directions, angular errors were standardized by reversing the sign for one of the rotations. In this convention, a positive error of 30 degrees indicates no compensation for the rotation, and values closer to zero reflect greater compensation. Complete compensation was defined as errors within ±3.3°, corresponding to the diameter of the ball. Ball trajectory distance was also recorded, and trials were excluded if the ball failed to travel at least 17 cm (20% of the required distance) or if angular errors exceeded ±60°. These criteria resulted in the removal of 96 trials (0.07% of all trials) due to distance and 586 trials (0.41%) due to excessive angular error.

To control for baseline tool or target biases, we computed the median angular error for each tool–target pair during the Baseline phase and subtracted these values from Training phase trials. For all statistical analyses, we focused on performance with the red paddle, which was the only tool common across all four experimental groups. This ensured comparability across the different cue conditions. While the second tool in each pairing was not included in the main analyses, performance for both tools is presented in Fig 3 for descriptive comparison. As expected, there were no substantial performance differences between tools during the Baseline phase or in the first block of Training trials prior to introducing the second tool and perturbation. Across groups, paddle performance (shown in purple in Fig 3) fell within the 95% confidence intervals of the other tool’s errors, suggesting no tool-specific performance biases at the start of adaptation.

**Figure 3.**
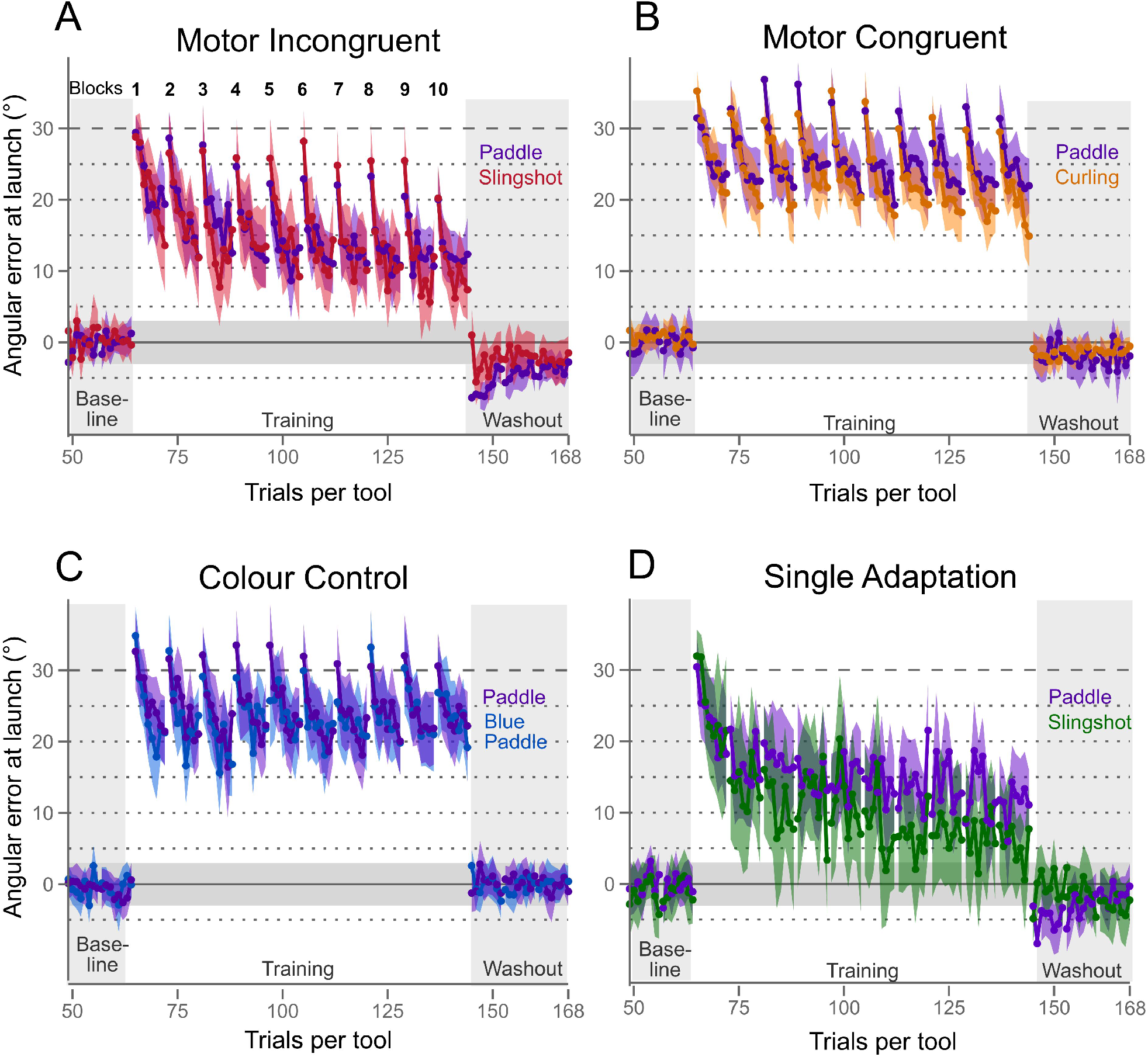
Launch angle error across tool trial numbers, averaged across participants for each tool pair, shown for the four groups: A) Motor Incongruent (purple for paddle, red for slingshot), B) Motor Congruent (purple for paddle, orange for curling rod), C) Colour-Control (purple for paddle, blue for blue paddle), and D) Single Adaptation (purple for paddle, green for slingshot). Although tools alternated in blocks of 8 trials, errors for both tools are plotted as overlapping across the corresponding trial number for each tool. These ten blocks are numbered on the top of panel A. The horizontal gray bar (±3° centered on the x-axis) indicates the range within which the 3.3° ball will hit the 3.3° target, relative to the launch point, for a successful hit. Shaded vertical areas on the left and right of each plot mark aligned trials, with the Baseline phase on the left and the Washout phase on the right. Only the final 24 trials of Baseline for each tool are displayed. The unshaded central area represents the Training phase, where ball motion was deviated 30-degree, with block numbers for this phase shown above panel A. Although the rotations applied to the ball were in opposite directions for the two tools, the sign of one rotation was reversed to facilitate comparison. 95% confidence intervals are shown in matching colour for the two tools of each group.

We observed a small but significant effect of tool order during training. Specifically, participants who encountered the paddle as the second tool showed larger mean errors (34.23°) on the first paddle trial than those who used the paddle first (27.98°), suggesting a 7° interference effect from prior exposure to the opposing perturbation.

This tool order effect was significant (F(1,134) = 7.478, p < 0.01), but it did not interact with group (F(3,134) = 0.67, p = 0.57). Because tool order and associated rotation direction were counterbalanced across participants, we collapsed across tool order in our primary analyses to simplify comparisons. While this approach may reduce sensitivity to detect between-group differences in learning rates due to increased variance, it provides a more balanced assessment of overall group performance across both tools, and our relatively large sample sizes help mitigate this reduced sensitivity.

To compare adaptation across the four groups, we conducted analyses at both the trial level and the block level using angular errors from the paddle. At the trial level, we compared the first and final trials of the Training phase to capture the full extent of individual trial-by-trial adaptation. At the block level, we averaged angular errors across all eight trials within each block and compared the first and final blocks of training. This dual-level approach allowed us to assess not only the immediate change in performance within a single trial but also broader patterns of learning across repeated exposures to the same tool–rotation pairing.

Statistically, we used a 4 (group: Congruent, Incongruent, Colour, Single) × 2 (time: early vs. late) linear mixed-effects model (LMER) separately for (1) the first and last trials of training and (2) the mean errors from the first and last blocks. This enabled us to examine overall group differences in the magnitude of adaptation, as well as any interactions indicating group-specific learning profiles. Where significant group × time interactions were found, we performed one-way ANOVAs on the first trial and first block to confirm that all groups started with comparable performance levels. As expected, there were no significant group differences in the first trial (F(3,138) = 0.43, p = 0.72) or first block (F(3,140) = 1.53, p = 0.21), supporting the assumption that all groups began training with similar baseline-corrected performance. These results were consistent even when the analysis was restricted to the three dual adaptation groups (all p > 0.05). We then compared performance in the final training block across groups using a one-way ANOVA to test whether any differences emerged over the course of training.

When significant effects were detected, Tukey post hoc comparisons were used to identify specific group differences. This stepwise approach ensured that any observed group differences in final performance could be interpreted as the result of learning under different cue conditions, rather than pre-existing performance differences or random variation in early trials.

To examine within- and between-block adaptation, we analyzed performance on the first trial (T1) and final trial (T8) of each 8-trial block. This allowed us to quantify both short-term learning and retention after exposure to the opposing perturbation. Although full T1 and T8 data are plotted across all blocks in Fig 4A–C, we focused our statistical comparisons on the first and final training blocks (Fig 3A). We used two 3 (group) × 2 (block) LMERs—one for T1 and one for T8—to assess learning and retention across the Training phase. Any significant group × block interactions were followed by one-way ANOVAs across groups on the first and final blocks to isolate the source of the interaction. Post hoc comparisons were conducted as needed.

**Figure 4.**
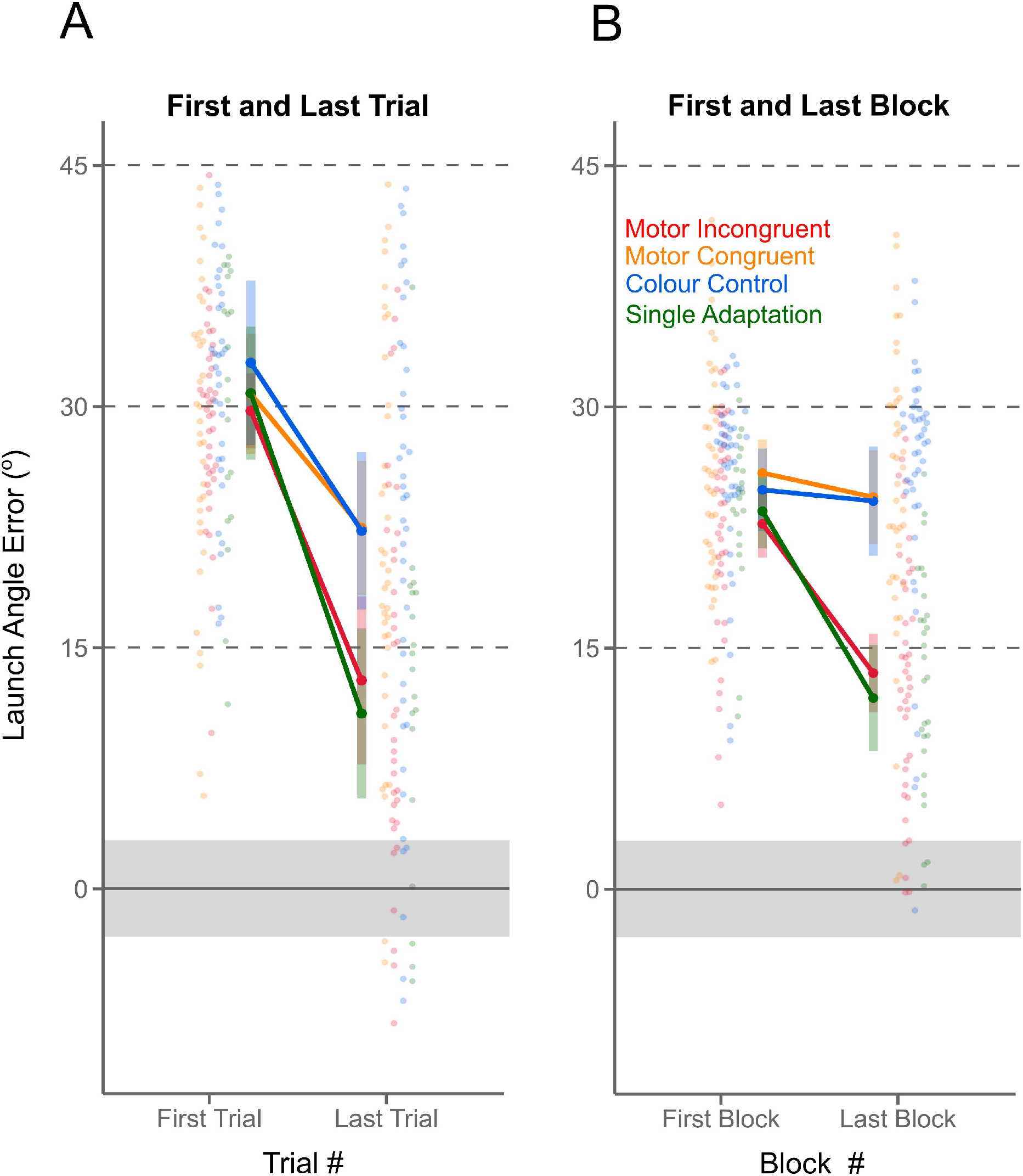
Angular launch error when adapting to the 30° rotation of ball direction using the paddle, shown for each participant (hollow circles) and for each of the four groups (full circle), with 95% confidence intervals across participants. (A) Launch errors averaged over the first and last trials of the Training phase. (B) Launch errors averaged over the first and last blocks of 8 trials during Training. Dark circles indicate the group mean error, with 95% confidence intervals across participants. The gray horizontal bar shows the range required for a successful hit, as illustrated in Figure 3.

To assess aftereffects and de-adaptation, we compared angular errors for the paddle between the final block of Baseline and the first block of Washout across all four groups using a 4 × 2 LMER. Significant interactions were followed by paired t-tests within each group to assess whether errors increased in the expected direction following removal of the perturbation. To control for multiple comparisons, a Bonferroni correction was applied to the four planned t-tests, resulting in an adjusted significance threshold of p = 0.0125.

Data and scripts can be found https://osf.io/khms8

Github: https://github.com/maceagle12/ToolUseData

## Results

To evaluate whether tool-related cues are sufficient for dual adaptation, we first describe the overall adaptation patterns across all tool pairings before focusing on performance with the common paddle tool. Each group used a distinct combination of two tools, with each tool consistently paired with a unique visuomotor perturbation. Our central aim was to determine whether adaptation to one tool would interfere with learning the opposing perturbation associated with the second tool—or whether certain contextual cues would allow participants to maintain separate motor memories and facilitate dual adaptation. To enable direct comparison across conditions, we focus our main statistical analyses on the red paddle, which was used in all four groups. By analyzing how performance with this shared tool changed in the presence of different paired tools, we could isolate the influence of contextual cue type—tool identity, control dynamics, or color—on adaptation.

### General Performance Patterns Across Tool Pairings

Figure 3 shows angular launch errors across the final three Baseline blocks, all ten Training blocks, and the first three Washout blocks for each group, plotted separately for each tool. To compare across opposing perturbation directions, errors from one tool in each pair were sign-reversed so that zero always reflects full compensation. Although tools were alternated in blocks of 8 trials, we plot their trial sequences as overlapping to visualize adaptation across the experiment.

As expected, all groups performed comparably at Baseline (all p > 0.18). Upon introduction of the visuomotor rotation, each group initially exhibited large deviations consistent with the imposed perturbation (first trial F(3,138) = 0.43, p = 0.72; first block F(3,140) = 1.53, p = 0.21) (First trial means: Motor Incongruent (m= 29.7, SE= 1.92),Motor Congruent (m= 31.2, SE= 1.92), Colour control (m= 32.7, SE = 1.85), Single adaptation (m = 30.8, SE = 2.72); First Block means: Motor Incongruent (m=22.7, SE = 1.11), Motor Congruent (m = 26.0, SE = 1.11), Colour Control (m = 24.9, SE = 1.05), Single Adaptation (m = 23.5, SE =1.56). All groups reduced their errors over successive trials, indicating adaptation. However, a distinctive zig-zag pattern emerged across Training for DA groups, reflecting interference: within-block improvements did not fully persist into the next block with the same tool, at least for the Motor Congruent and Control Control group, suggesting the intervening tool disrupted retention.

The Motor Incongruent group also showed a zig-zag pattern, but with clearer evidence of retention across successive blocks using the same tool, suggesting reduced interference compared to the other dual-adaptation groups. First-trial errors decreased across successive paddle blocks, suggesting greater retention across tool switches.

This pattern was less apparent in the Colour and Motor Congruent groups, where within-block learning was not maintained between blocks. These differences are analyzed in more detail below.

### Dual Adaptation: Impact of Tool Pairing

To quantify group differences in adaptation, we examined paddle performance during Training. There was a significant main effect of trial, with angular error decreasing from the first to the final trial (F(1,138) = 79.31, p < 0.001; Fig. 4A), and a significant main effect of block, with angular error decreasing from the first to the final block (F(1,147) = 61.02, p < 0.001; Fig. 4B). However, the magnitude of this change differed across groups, as indicated by significant trial × group and block × group interactions (trial × group: F(3,139) = 2.84, p < 0.05; block × group: F(3,148) = 13.33, p < 0.001). Post hoc tests revealed that only the Motor Incongruent group (red) did not differ significantly from the Single Adaptation group (green) by the end of Training (p = 0.928) as illustrated in green in Fig 4. In contrast, both the Colour (blue) and Congruent (orange) groups showed significantly less improvement (both p < 0.001), with no difference between them (p =0.999). These results indicate that only Motor Incongruent tool pairings enabled sufficient contextual separation to support dual adaptation.

### Retention Across Alternating Blocks

To investigate how adaptation was retained across tool switches, we examined performance across the ten alternating training blocks. Specifically, we plotted launch errors for the first trial (T1) of each block (Fig 5A) and the last trial (T8) of each block (Fig5C). This allowed us to assess, respectively, how much adaptation was retained at the beginning of each block and how much cumulative learning was achieved within each block over time. Visual inspection of Fig 5A shows a progressive reduction in initial error across blocks for the Motor Incongruent group (red), indicating that participants retained some adaptation from previous encounters with the same tool—even after using the other, differently perturbed tool in the intervening block. In contrast, the Motor Congruent (orange) and Colour Control (blue) groups show relatively flat trajectories across blocks, with little reduction in initial error, suggesting weaker retention across tool switches.

**Figure 5.**
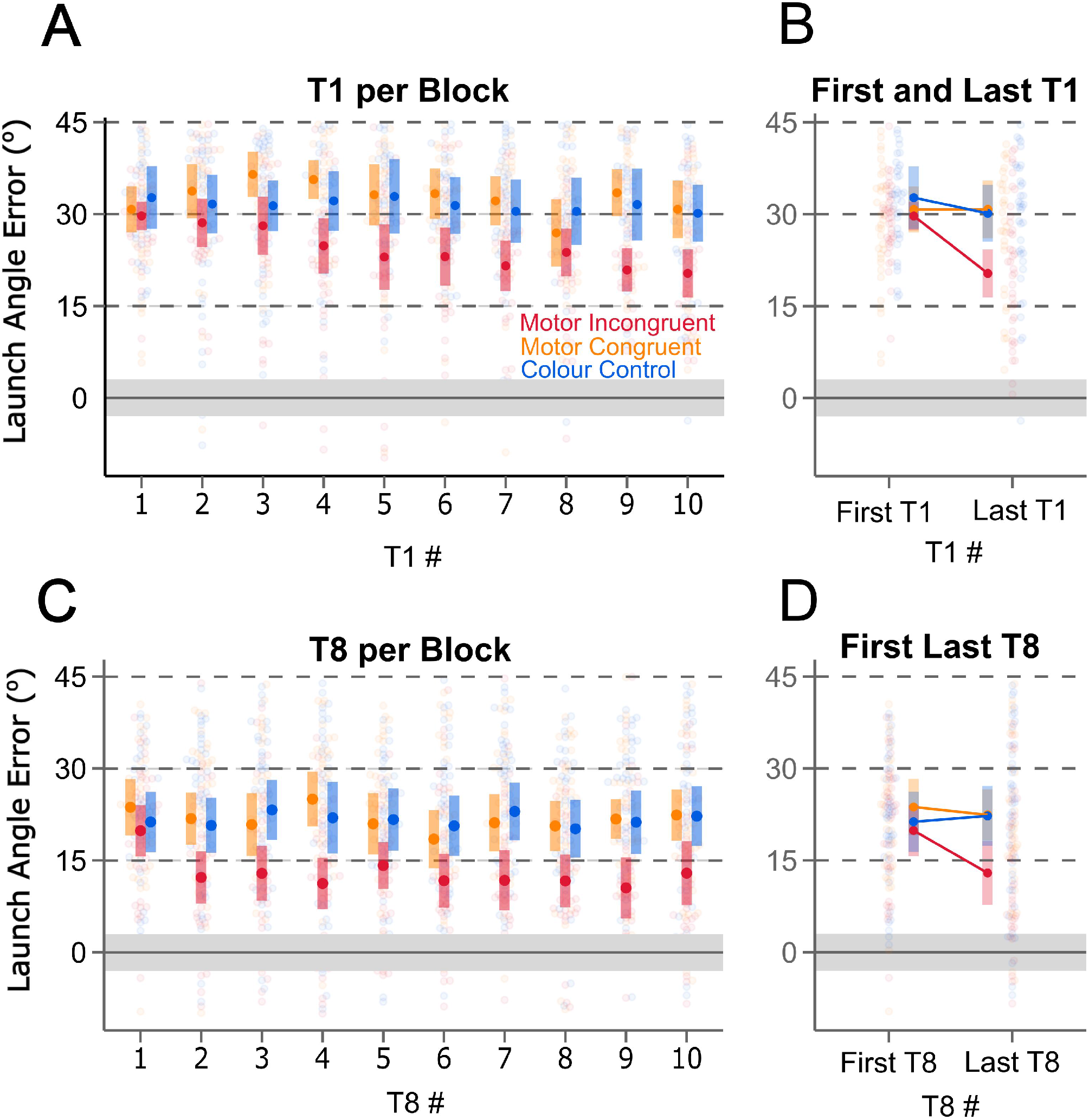
Launch errors for first trial (T1) (top row) and final trial (T8) (bottom row) of every block (A, C) and the first and last block (B, D) for each participant (hollow circles), and averaged across participants (solid circles, with bars representing 95% confidence interval) for the three Dual adaptation groups; Motor Incongruent (red), Motor Congruent (orange), Colour Control (blue). Grey horizontal bar as in Figure 3.

In Fig 5C, which depicts the final trial of each block, a similar—but less pronounced— trend is observed for the Motor Incongruent group: errors decrease modestly over the first few blocks. In contrast, the other two groups show no change across all 10 blocks. However, there is greater variability in the end-of-block data compared to the first trials, and the downward trend is less pronounced.

To quantify these retention effects, we focused on comparing only the first and last blocks of training (Blocks 1 and 10) for both T1 and T8 trials in the Dual Adaptation conditions—where the greatest change in performance would be expected due to repeated exposure (Figs 5B and 5D). Comparing launch error for T1 in the initial and final blocks (Fig 5B) reveals a significant reduction in error from Block 1 to Block 10 (F(1,120) = 7.12, p < 0.05). This confirms that some adaptation achieved within each block is retained in subsequent blocks, for the same tool. However, this retention was group-dependent, as revealed by a significant group × block interaction for T1 (F(1,120) = 3.45, p < 0.05). Because the groups did not differ in their T1 performance on the first block (as previously reported), this interaction reflects differences in performance during the final block. A follow-up ANOVA on the final block confirmed a main effect of group on T1 (F(2,119) = 6.87, p < 0.01). Tukey comparisons showed that the Motor Incongruent group retained more learning across blocks than both the Colour Control (p < 0.01) and Motor Congruent groups (p < 0.01). In fact, only the Motor Incongruent group shows a visible reduction in error from Block 1 to Block 10: the red data point in Fig 5B falls below the confidence interval of their initial value.

When comparing T8 trials across blocks (Fig 5D), no significant change in error was detected (F(1,120) = 1.96, p = 0.16), and no group × block interaction was found (F(1,120) = 1.84, p = 0.16). This suggests that the cumulative adaptation achieved by the end of the first block (before any interference from the second tool) was not substantially improved after additional practice. This lack of statistical significance may reflect increased inter-subject variability, rather than the absence of a true effect. particularly in the Motor Incongruent group where the mean trend did suggest a reduction in error.

Overall, these findings suggest that tool-pairings involving Motor Incongruent motor actions, which provided both intrinsic and extrinsic cues, supported greater retention of context-specific learning across alternating blocks. The Motor Incongruent group’s performance—particularly in the first trial of each block—shows evidence of dual adaptation that accumulates across training despite frequent switching between tools with opposing perturbations.

### Aftereffects Following Adaptation

To evaluate persistent adaptation, we compared angular errors in the first Washout block to the final Baseline block, as illustrated in Fig 6. A significant group × phase interaction (F(3,140) = 20.60, p < 0.001) indicated that aftereffects differed by condition. Only the Motor Incongruent (red) and Single Adaptation (green) groups showed significant deviations during Washout (both p < 0.001, Bonferroni corrected), consistent with implicit motor memory formation. The Congruent and Colour groups showed no such aftereffects (both p > 0.0125), reinforcing the conclusion that visual or identity cues alone were insufficient to support robust durable dual adaptation.

**Figure 6.**
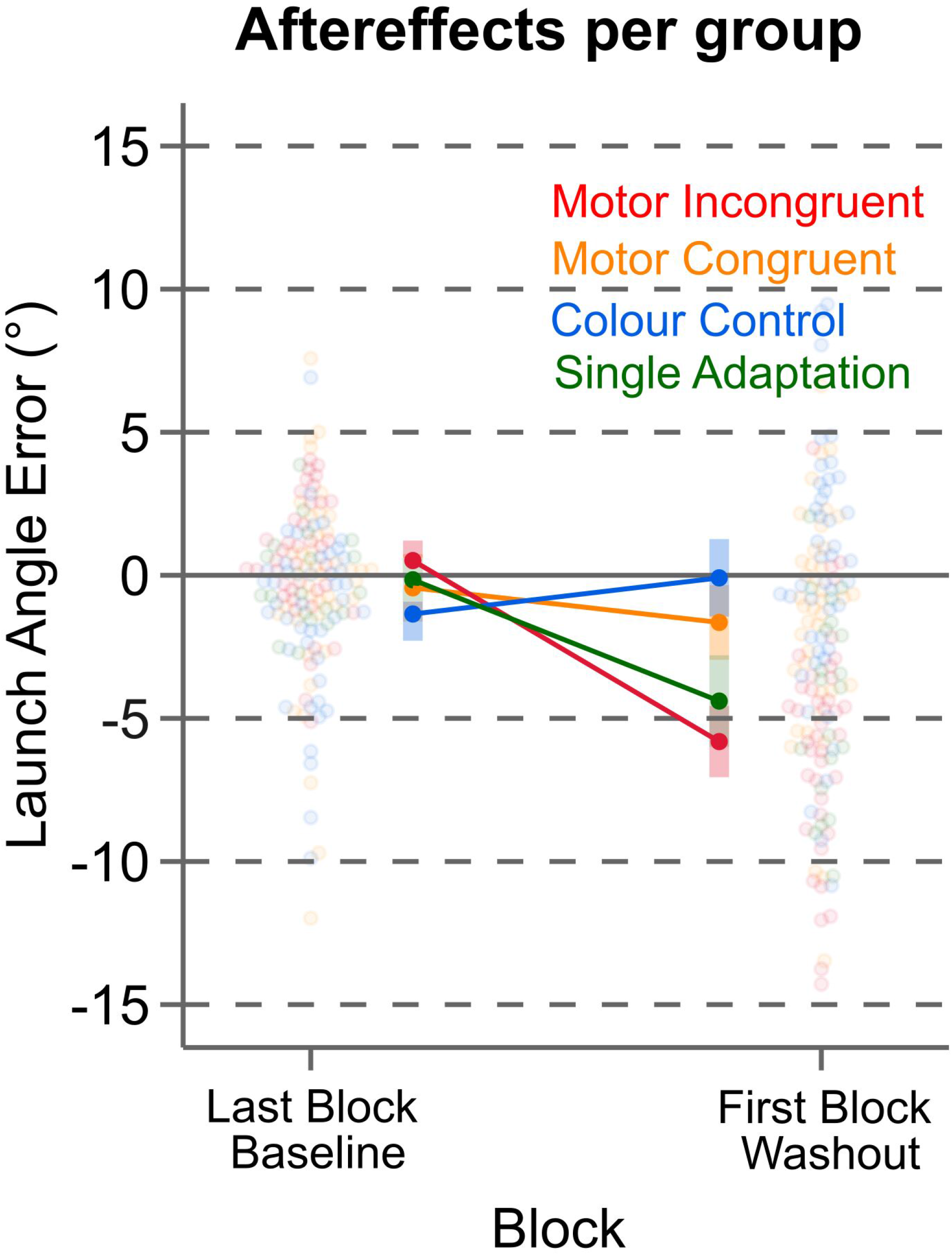
Mean launch error (filled circles) for paddle across all four groups across the Last block of Baseline and the First Block of Washout. Individual participant error (hollow circles) and 95% CIs (coloured rectangles) are shown for all trials.

## Discussion

The present study demonstrates that distinct tool operations—not visual features alone—enable successful dual adaptation to opposing visuomotor perturbations in immersive VR. When tools differed in both appearance and how they were operated (Motor Incongruent group), participants adapted as effectively as in the Single Adaptation group, despite alternating between tools with opposing perturbations every eight trials. In contrast, when tools differed only in color (Colour group) or had distinct visual appearances but shared similar tool wielding patterns (Motor Congruent group), participants exhibited limited retention across blocks and failed to show aftereffects— suggesting substantial interference and a lack of distinct motor memory formation. These findings build on prior work emphasizing the importance of intrinsic contextual cues—those related to how a tool is operated—in enabling dual adaptation (e.g., Ayala et al., 2015b; Heald et al., 2018, 2021; Howard et al., 2015; Proud et al., 2019). Our results further support the conclusion that extrinsic cues are insufficient unless tightly coupled with distinct control policies ^5–8,11–14,20,21,29,30^. Previous studies testing extrinsic cues typically relied on minimal differences—such as changes in color or shape—that were not directly relevant to the task. In contrast, our study is one of the first to employ rich, task-relevant extrinsic cues: the paddle and shuffle tools were visually distinct and differed slightly in their method of operation. Despite this, our findings show that even these salient visual distinctions, in the absence of clearly differentiated tool-use dynamics, do not support robust dual adaptation.

Notably, adaptation in the Motor Incongruent group was characterized by progressive improvement in the first trial of each block of the same tool, suggesting increasing retention despite blocks of interleaved perturbations linked with the second tool. This pattern was absent in the Colour and Congruent groups, where adaptation had to be re-learned in each block. These results align with the notion that interference in dual adaptation arises when the same control policy is reused across contexts ^30^, and that differences in tool operation help the motor system retrieve and retain the appropriate internal model.

Our aftereffects findings further underscore the importance of intrinsic cues related to tool operation. In both the Motor Incongruent and Single Adaptation conditions, participants exhibited significant reach deviations when the visuomotor perturbation was removed for the common paddle—indicating the formation of implicit motor memories. In contrast, no aftereffects were observed in the Colour Control or Motor Congruent groups, where limited separation between internal models likely prevented the consolidation of implicit adaptation.

Interestingly, the slingshot tool—despite supporting effective adaptation during training—did not produce observable aftereffects in either the Dual Adaptation or Single Adaptation conditions (see Fig 6). This discrepancy may reflect that the slingshot was somewhat easier to learn, or alternatively, that participants relied on explicit strategies due to its constrained and repetitive draw-and-release action. The uniformity of this movement pattern may have encouraged the use of simple aiming heuristics or rule-based strategies, thereby reducing reliance on an internal forward model ^31,32^. This may also account for the slightly greater compensation observed during training with the slingshot, although the performance advantage over the paddle was modest— approximately 5–10%. In contrast, the paddle, which permitted a wider range of movement directions and magnitudes, likely required more flexible motor recalibration and thus engaged implicit learning mechanisms that persisted even after the perturbation was removed.

This study extends dual adaptation research by explicitly manipulating tool dynamics within an immersive VR environment, where spatial cues and the surrounding visual context were held constant. The use of VR provided a unique advantage by allowing us to control for environmental variables while systematically testing both tool-specific dynamics and richer, task-relevant extrinsic cues, such as distinct visual tool identities (e.g., Modchalingam et al., 2025). Our findings more conclusively show that even these salient visual features (like tool identity) —despite being relevant to the task—are not sufficient to support dual adaptation in the absence of distinct operational demands.

Unlike prior studies that relied on workspace separation ^14,15,17,22,33,34^), explicit lead-in movements (^6–8,11,21^, or movement planning manipulations (^30^), we demonstrate that tool-specific operational demands alone can serve as effective contextual signals. Our design represents an important step toward mimicking naturalistic tool use and supports the idea that tools are encoded primarily by their functional demands rather than their visual appearance ^35–39^.

The COIN model ^28,40^ offers a useful framework for interpreting our results by proposing that the motor system forms and retrieves internal models based on probabilistic inference about the current context. Rather than switching between fixed motor memories, the brain continuously evaluates sensory cues and prior experience to estimate which motor memory—or combination of memories—is appropriate at a given moment. In our study, only the Motor Incongruent group demonstrated clear evidence of dual adaptation, consistent with the idea that distinct tool operations for launching the ball provided strong, unambiguous cues to signal different contexts. In contrast, tool identity alone (as in the Color Control and Motor Congruent groups) may have lacked the contextual specificity required to support reliable contextual inference, resulting in interference. It remains possible that with extended training, these extrinsic cue conditions might eventually support the formation of separate motor memories; however, blocks of eight training trials per visual tool appeared insufficient. Overall, our findings align with the COIN model’s central claim that the reliability and task-relevance of contextual cues are critical determinants of whether distinct internal models are formed and retrieved.

Our findings also suggest that the effectiveness of contextual cues in supporting dual adaptation may not stem solely from prompting the formation of entirely distinct internal models, but rather from reducing interference from the competing memory—thereby enhancing retention across blocks with the same cue. As shown in Figures 3 and 4, the Motor Incongruent group exhibited increasingly stable performance with each return to a given tool, as interference between the opposing mappings diminished with additional training. Notably, this did not lead to greater learning *within* each block—at least not in a clearly observable way. This suggests that the cues or contexts did not facilitate faster or more complete formation of separate motor memories during each exposure, but instead supported gradual strengthening and maintenance of those memories through repeated retrieval with the same cue. From this perspective, dual adaptation may sometimes reflect improved protection of a motor memory in the face of an opposing perturbation, rather than the creation of new internal models, underscoring the role of cue distinctiveness in limiting interference between competing mappings.

This also suggests that paradigms in which cues for different perturbations are presented in a randomly interleaved fashion—without a minimum number of consecutive trials per context—may fail to elicit dual adaptation not because the cues are inherently ineffective, but because they do not allow sufficient time to establish a stable memory trace for each condition. When exposure to a given cue-perturbation pairing is too brief or interrupted too frequently, the motor system may be unable to reinforce the association strongly enough to support retention. Mini-blocked designs, like the one used here and by others who demonstrate dual adaptation^14,16,28,40–42^, may be especially important for scaffolding the initial stabilization of multiple memories, even if eventual flexible switching is the end goal. Concurrent adaptation to opposing perturbations that alternate randomly typically results in weaker dual adaptation, unless the contextual cues are highly salient or strongly associated with each perturbation ^11,12,15,22,24^. This highlights the importance of considering not just the nature of contextual cues, but also the structure and temporal continuity of exposure when designing dual adaptation paradigms.

In conclusion, our results highlight the importance of movement-relevant and operational cues for organizing multiple motor memories. The findings have practical implications for tool-based training and rehabilitation, suggesting that designing tools with distinct control demands may help reduce interference and enhance learning in complex motor tasks.

